# Subgenomic RNAs as molecular indicators of asymptomatic SARS-CoV-2 infection

**DOI:** 10.1101/2021.02.06.430041

**Authors:** Chee Hong Wong, Chew Yee Ngan, Rachel L. Goldfeder, Jennifer Idol, Chris Kuhlberg, Rahul Maurya, Kevin Kelly, Gregory Omerza, Nicholas Renzette, Francine De Abreu, Lei Li, Frederick A. Browne, Edison T. Liu, Chia-Lin Wei

## Abstract

In coronaviridae such as SARS-CoV-2, subgenomic RNAs (sgRNA) are replicative intermediates, therefore, their abundance and structures could infer viral replication activity and severity of host infection. Here, we systematically characterized the sgRNA expression and their structural variation in 81 clinical specimens collected from symptomatic and asymptomatic individuals with a goal of assessing viral genomic signatures of disease severity. We demonstrated the highly coordinated and consistent expression of sgRNAs from individuals with robust infections that results in symptoms, and found their expression is significantly repressed in the asymptomatic infections, indicating that the ratio of sgRNAs to genomic RNA (sgRNA/gRNA) is highly correlated with the severity of the disease. Using long read sequencing technologies to characterize full-length sgRNA structures, we also observed widespread deletions in viral RNAs, and identified unique sets of deletions preferentially found primarily in symptomatic individuals, with many likely to confer changes in SARS-CoV-2 virulence and host responses. Furthermore, based on the sgRNA structures, the frequently occurred structural variants in SARS-CoV-2 genomes serves as a mechanism to further induce SARS-CoV-2 proteome complexity. Taken together, our results show that differential sgRNA expression and structural mutational burden both appear to be correlated with the clinical severity of SARS-CoV-2 infection. Longitudinally monitoring sgRNA expression and structural diversity could further guide treatment responses, testing strategies, and vaccine development.

## Introduction

COVID-19, emerged in late 2019, was caused by severe acute respiratory syndrome coronavirus 2 (SARS-CoV-2). With its high infectivity and mortality rates, particularly in individuals of older age and those with pre-existing health conditions, COVID-19 has rapidly expanded into a global pandemic. Of great importance in the management of the pandemic is the observation that many infected individuals are asymptomatic, ranging from 20-80% ^1,2,3^. Asymptomatic patients, while having faster viral clearance ^4–7^, appear to have similar viral loads compared to symptomatic patients ^4,5,8–11^ and, therefore, can effectively transmit the disease. Since viral load is not a reliable predictor of disease severity, we examined the genomic biology of SARS-CoV2 infection in primary patient samples for other correlates of clinical severity.

To understand the pathophysiology of COVID-19 infection, major effort has been made on fully decoding the SARS-CoV-2 genome and its genetic variation, specifically the single nucleotide variants (SNVs) ^12,13,14^. SARS-CoV-2 is a positive, single-stranded RNA virus. Upon infecting into the host cells, the viruses deploy both replication and transcription to produce full-length genomic ~30-Kb RNAs (gRNAs) and a distinct set of “spliced” subgenomic transcripts (sgRNAs). These sgRNAs are transcribed through a “discontinuous transcription” mechanism ^15^ by which negative-strand RNAs are produced from the 3’ of gRNAs followed by a template switch from a 6-nucleotide ACGAAC core transcription regulatory sequence (TRS) that are complementary between 5’ TRS-Leader (TRS-L) and a set of individual TRS-Body (TRS-B) at the 3’-end of the viral genome to join with individual open reading frames (ORFs). These distinct sgRNAs are subsequently served as viral mRNAs for translation of multiple structural and accessory proteins including spike surface glycoprotein (S), small envelope protein (E), matrix protein (M), and nucleocapsid protein (N) ^16^. SgRNAs are not packaged into virions and only transcribed in infected cells, therefore, their presence is thought to be an indicator of effective viral replication and *in vivo* host fitness ^17–19^.

So far, only a few studies have examined their sgRNAs and primarily were investigated in *in vitro* cell culture models ^20–22^. Despite emerging evidence showing the impact of structural variants in the sgRNA coding regions on the severity of infection, transmission rates and immune responses ^23,24^, SARS-CoV-2 structural variants and sgRNAs, particularly their abundance and complexity in the context host response have been grossly overlooked. We posited that the molecular characterization of the SARS-CoV-2 associated with asymptomatic infection could help to understand virulence factors contributing to viral pathogenicity and regulation of host responses.

Here, we systematically characterized the diversity and prevalence of structural deletions and sgRNA expression in primary human tissues from both symptomatic and asymptomatic individuals using a suite of genomic and transcriptomic analyses. From routine swabs collected for diagnostic purpose, we ascertained sgRNA configurations and found that their abundance, both as individual sgRNA species and collectively as a group, is drastically reduced in asymptomatic infection. Moreover, we identified widespread structural deletions in the SARS-CoV-2 genomes, particularly in the regions encoding sgRNAs. Distinct sets of deletions can be consistently and preferentially found in independent SARS-CoV-2 genomes associated with symptomatic and asymptomatic cases, respectively, suggesting their functional significance. To understand the impact of structural variants on the viral protein integrity, we analyzed the predicted viral proteomes from full-length viral transcript isoforms. Our results reveal the highly unstable nature of SARS-CoV-2 genomes and reveal the potential utility of sgRNA expression as an indicator of clinical severity.

## Results

### SgRNA expression is drastically repressed in asymptomatic SARS-CoV-2 infection

SARS-CoV-2 gRNAs and sgRNAs have overall high sequence identity. To discern sgRNA from the gRNAs, we exploited the features derived from the discontinuous transcription, namely the joining between TRS-L and TRS-B regions whose presence exclusively was found in sgRNAs. We adopted amplicon-based sequencing (amplicon-seq), a method widely used to characterize SARS-CoV-2 genomes ^25^, to characterize the presence of sgRNAs and correlate their abundance in the COVID-19 positive samples between symptomatic and asymptomatic patients. Amplicon-seq is highly sensitive, with limit of detection (LoD) reported as low as one SARS-CoV-2 copy per microliter using the optimized protocols from the Artic network ^26,27^. Therefore, it can effectively enrich for SARS-CoV-2 cDNAs from samples of wide-range of viral content. In this approach, viral-specific primers were designed across the full length RNAs and amplicons specific for SARS-CoV-2 sgRNAs can be PCR amplified by 5’ most primer next to the TRS-L sequence as forward primer and reverse primers nearest to the TRS-B sequences in the multiplex PCRs. Based on the locations of primers, we anticipated that amplicons for 6 out of the 9 sgRNA species (sgRNA_S, E, M, 6, 7b and N) can be found in the amplicon-seq (Methods). Followed by massively parallel sequencing, these subgenomic-specific amplicons can be identified through the junction reads linking TRS-L and TRS-B in the sequencing data and used to determine the relative abundance of sgRNAs (Fig. 1a).

**Figure 1:**
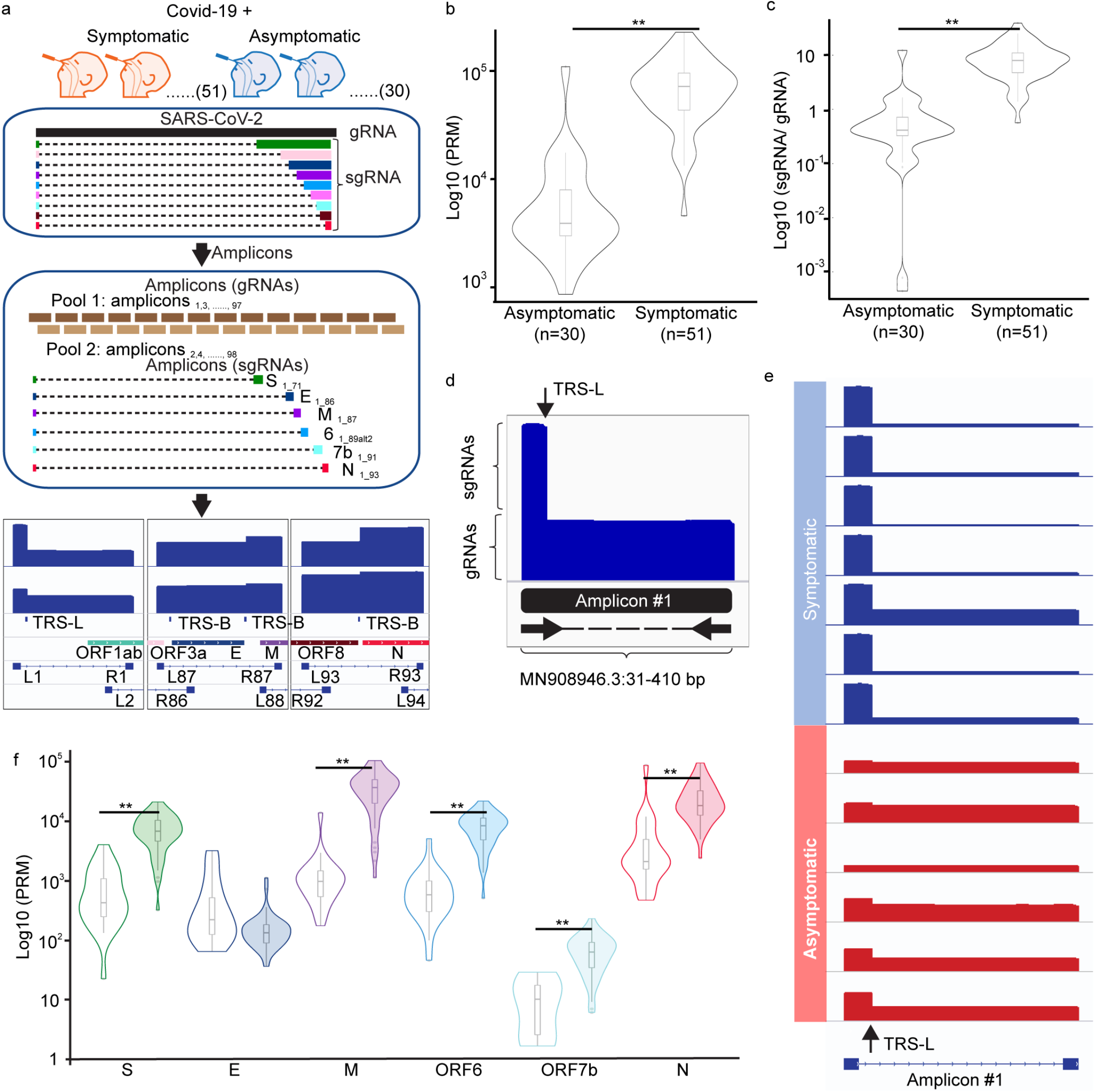
Amplicon-seq analysis characterize sgRNA relative abundance in clinical swabs in COVID-19 positive patients. **a.** SARS-CoV-2 RNAs existed in the upper respiratory tracts from both symptomatic and asymptomatic COVID-19 positive patients were analyzed by Amplicon-seq. Amplicons tilted across full-length SARS-CoV-2 genomes were amplified by specific primers in the multiplex PCRs in two separate pools. Subgenomic RNAs (sgRNAs) specific amplicons can be identified by specific amplification with 5’ forward primer closest to the transcription regulatory sequence leader (TRS-L) site and 3’ reverse primers closest to the TRS-Body (TRS-B) sites followed by sequencing and alignment. Distribution of sgRNAs normalized counts **(b)** and sgRNA to genomic RNA (gRNA) ratio (*p*=4.9 × 10^−12^) **(c)** between symptomatic and asymptomatic cases (*p* = 5.6 × 10^−12^). **d.** Sequencing coverage across the 5’ 400 nucleotides of SARS-CoV-2 genome shows the contribution from sgRNAs and gRNA, respectively. **e.** Amplicon-seq coverages across 5’ 400 nucleotides from representative symptomatic and asymptomatic cases. **f.** Distribution of the normalized counts of individual sgRNA species measured in symptomatic and asymptomatic cases (*P* values of pair-wise comparison for S, M, ORF6, ORF7b and N are 6 × 10^−11^, 9 × 10^−12^, 2 × 10^−11^, 2 × 10^−7^, and 9 × 10^−10^). All statistical tests are two-sided Wilcoxon Rank-Sum Test. Center line, median; boxes, first and third quartiles; whiskers, 1.5 × the interquartile range.

From 51 and 30 SARS-CoV-2 positive symptomatic and asymptomatic patients respectively (where asymptomatic patients are defined as those who showed none of the key COVID-19 symptoms within 14 days of testing) (Supplementary Table 1), we extracted total RNA from swabs of different locations of respiratory tracts including anterior nasal, oro- and naso-pharyngeal collected for the purpose of diagnostic RT-PCR and performed amplicon-seq to generate deep sequencing data for each sample (>200,000 paired reads, >4000-fold genome coverage) (Supplementary Table 2). From the reads aligned to the reference MN908947.3, amplicon corresponding to 6 sgRNAs were detected through split-mapped reads connecting the first 75 nucleotides harboring TRS-L sequences to their respective TRS-B sites. To evaluate their relative abundance among different samples, we normalized the amounts of TRS-L associated junction reads against total numbers of SARS-CoV-2 reads in each sample. Through the normalized junction read counts, we found that the levels of sgRNAs were highly variable, ranging between 0 to 230,155 reads per million (RPM). Between COVID-19 positive individuals with and without symptoms, sgRNA levels were significantly lower in asymptomatic than in the symptomatic samples (median value 3,498 vs. 72,231; two-sided Wilcoxon Rank-Sum Test, *p* = 4.9 × 10^−12^) (Fig. 1b). To ensure that the reduction of sgRNA expression was not resulted from potential lower viral load found in the asymptomatic samples, we further compared the expression of sgRNA per viral gRNA (sgRNA/gRNA) in the asymptomatic *vs.* symptomatic infections. Here, the levels of gRNAs were defined as the amount of reads aligned uninterrupted across the first 400 nucleotides because their existence was exclusively found in the viral gRNA molecules. As shown in Fig. 1c, significantly lower ratio of sgRNA/gRNA (19-fold in median value, two-sided Wilcoxon Rank-Sum Test, *p* = 5.6 × 10^−12^) was observed in asymptomatic hosts, suggesting the lower levels of sgRNAs were independent of virus quantity in these samples. The relative abundance of sgRNAs to gRNAs can also be reflected through the read coverage along the first 400 nucleotides (Fig. 1d). Here, the distinct differences of sgRNA/gRNA ratio can be observed by the apparent degrees of differential coverage from the first 75 nucleotides (present in both sgRNAs and gRNAs) to the 76-400 nucleotides (only present in gRNAs) visualized through the Integrated Genomics Viewer (IGV) (Fig. 1e), which clearly indicated the existence of higher amount of sgRNAs in the symptomatic samples.

To evaluate if the reduction of sgRNAs occurred selectively in specific sgRNA species or broadly to all sgRNA transcription, we further compared the levels of each gRNA species detected between symptomatic *vs.* asymptomatic samples. The expression levels of individual sgRNA species were determined by assigning each TRS-associated junction read to their respective sgRNA origins based on their corresponding TRS-B site usage. Among the 6 sgRNA-specific amplicons produced in the amplicon-seq, all but one (sgRNA_E) displayed significant reduction (two-sided Wilcoxon Rank-Sum Tests, *p*-values 2 × 10^−7^ to 9 × 10^−12^) (Fig. 1f). Among them, sgRNA_M exhibited the highest degrees (6-37 fold) of decline. Collectively, these results indicated that the lack of active viral transcription in the asymptomatic infection and the sgRNA to gRNA ratio in the host cells appears to reflect the degree of disease severity.

### Coordinated expression of sgRNAs in primary human cells of symptomatic infection

The differential sgRNA abundance detected in COVID-19 positive samples between symptomatic and asymptomatic patients implicates their potential function in eliciting host responses. To characterize their expression in the infected cells of symptomatic patients, we adopted an unbiased metagenomic RNA-seq approach to survey the types of sgRNAs expressed and quantitatively evaluate their relative abundance in these samples (Supplementary Table 3). In metagenomic RNA-seq analysis, both host and SARS-CoV-2 RNAs expressed were comprehensively revealed by the sequencing of the extracted total RNAs. Using the centrifuge algorithm ^28^, we conducted full metagenome profiling and taxonomy classification to assess their relative ratio between human and SARS-CoV-2. Despite their relative low *Ct* values (13-19), suggesting of high viral content, the ratio of reads aligned to SARS-Cov-2 were highly variable among these samples, ranging from 0.06% to 78% (Fig. 2a).

**Figure 2:**
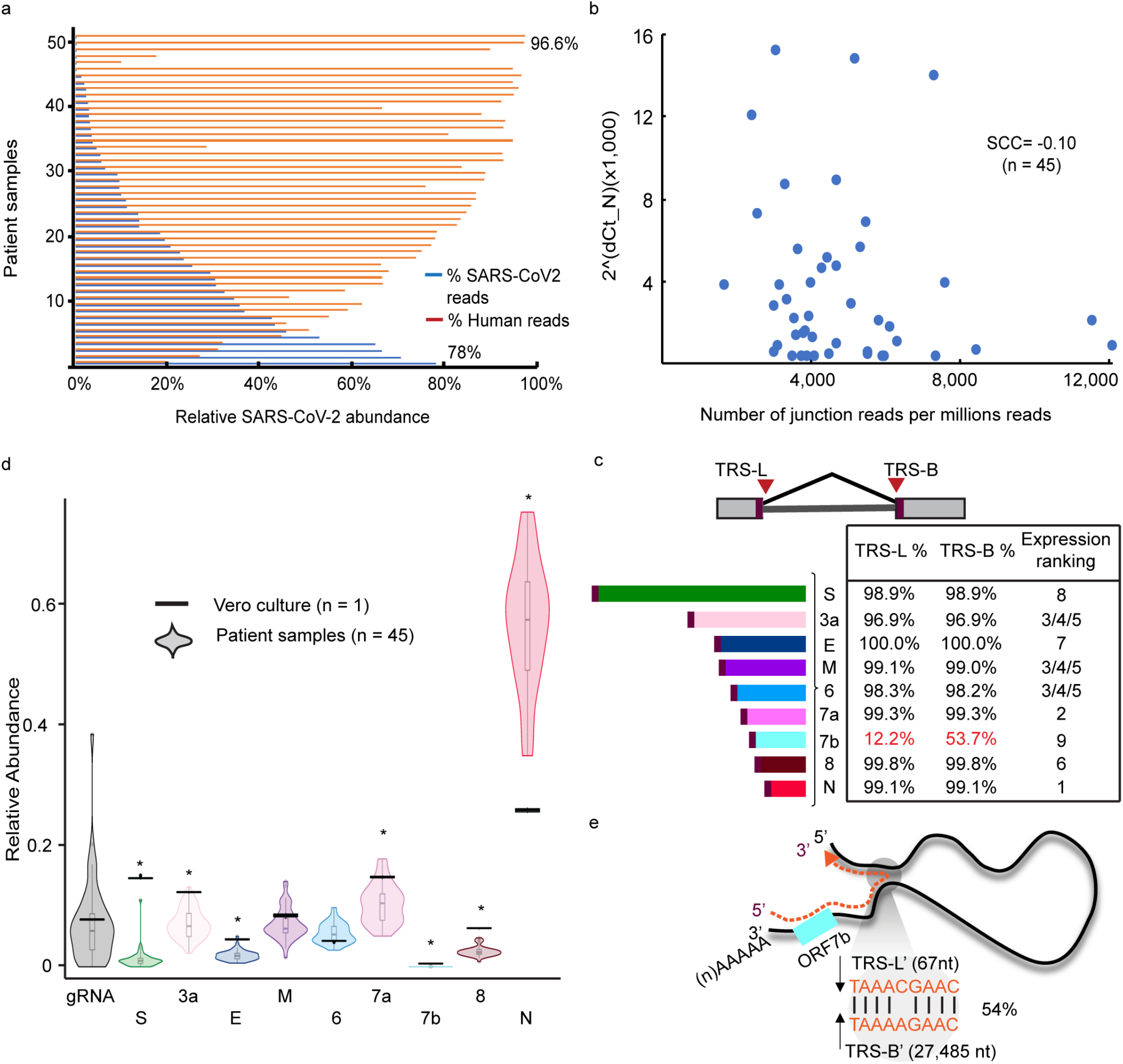
Expression of subgenomic RNAs (sgRNAs) in the clinical specimens from symptomatic patients. **a.** The percentages of SARS-CoV-2 (blue) and human (red) reads detected in each of the symptomatic samples (n=51). **b.** Correlation analysis between viral load (RT-qPCR *Ct* values) and sgRNA abundance (numbers of junction reads per million). **c.** Transcription regulatory sequence (TRS) usage. Percentages of sgRNA-derived junction reads split at their corresponding known TRS-Leader (TRS-L) and TRS-Body (TRS-B) sites for each sgRNA species and the relative abundance ranking. **d.** Proportions of reads assigned to genomic RNA (gRNA) and each sgRNA species in symptomatic samples (n=45) and Vero cultured cells (n=1). Center line, median; boxes, first and third quartiles; whiskers, 1.5 × the interquartile range; points, outliers. **e.** Sequences at the alternative TRS-B sites used by sgRNA_ORF7b transcription.

We next characterized the types and abundance of sgRNAs expressed in these samples. The TRS-L associated RNA-seq reads were assigned to each of the nine distinct sgRNA species based on their spans across the corresponding TRS-B junction sites closest to the annotated transcript initiation sites. The abundance of SARS-CoV-2 sgRNAs had no correlation with the viral load inferred by the *Ct* values from RT-qPCR testing (Spearman correlation coefficient = −0.10, *p* = 0.50) (Fig. 2b), suggesting that the viral nucleic acid shedding measured by the RT-qPCR diagnostic assays does not reflect the activity of viral replication in these samples. The relative abundance of different sgRNA species exhibited a remarkable consistency both in their expression ranking (Fig. 2c) and the relative proportion of the reads for each sgRNA class (Fig. 2d). Across all samples. SgRNA_N was expressed the highest and sgRNA_ORF7b was the least abundant. It is worth noting that sgRNA_ORF7b was not previously detected in *in vitro* infected cell cultures ^22^. The low expression of sgRNA_ORF7b could be resulted from the imprecision of TRS usage in the discontinuous transcription process. Unlike the other sgRNA species which were mostly transcribed from the annotated TRS sites, 54% of the sgRNA_7b transcripts have adopted an alternative TRS-B’ site MN908947.3:27485 (Fig. 2e). These observations suggested that ORF7b expression is subjected to high variability and could be dispensable *in vivo*.

When comparing the relative abundance of sgRNAs to these reported from *in vitro* Vero cells experiments, 7 sgRNAs exhibited significant difference (*p*-value < 1e-05) with the most striking difference found in the sgRNA_Spike (S) (Fig. 2d). In primary human samples, sgRNA_S expressed at less than 1% of total sgRNAs but was found at 14% of total expressed sgRNAs in the cultured Vero cells. The difference could be contributed by the differences in SARS-CoV-2 transmission and entry between the *in vitro* cell cultures and primary tissues. The expression of sgRNA_ORF10 was not detected, consistent with what has been described in SARS-CoV-2 infected cell cultures ^21^.

### Distinct sets of deletions detected in SARS-CoV-2 RNAs from primary human cells between symptomatic and asymptomatic infections

It has been reported that novel deletions in sgRNAs may have an impact on the clinical presentation of SARS-CoV-2 infection ^23^ and transmission rate ^29,30^. We therefore examined the structural deletions in SARS-CoV-2 RNAs found within symptomatic and asymptomatic individuals. Through the split-aligned reads that were not mediated from the TRS sites in the amplicon-seq data, we detected up to 10^4^ per million of SARS-CoV-2 paired reads harboring TRS-independent junctions of at least 20 nucleotides in each sample. These deletion events were more prevalent in viral samples from symptomatic hosts (two-sided Wilcoxon Rank-Sum Test, *p* = 2.3 × 10^−8^) (Fig. 3a), potentially due to more active viral replication in these hosts resulting in greater production of structural variants. In total, we detected 8,551 unique deletions in viral RNAs that were supported by ≥ 2 independent reads. While vast majority of them were sporadic events occurred in isolated cases, 501 (6%) deletions were consistently observed in >10% of samples; either specific in symptomatic (n=375), asymptomatic hosts (n=38) or in both (n=88) (Fig. 3b). It is interesting to note that, in symptomatic cases, these frequent structural deletions were not only more abundant but also significantly larger in sizes (median spans 198 vs 46 nucleotides, *p* = 1.6 × 10^−15^), pointing to a potential selection force for different types of viral variants adapted in distinct cohorts of host responses.

**Figure 3:**
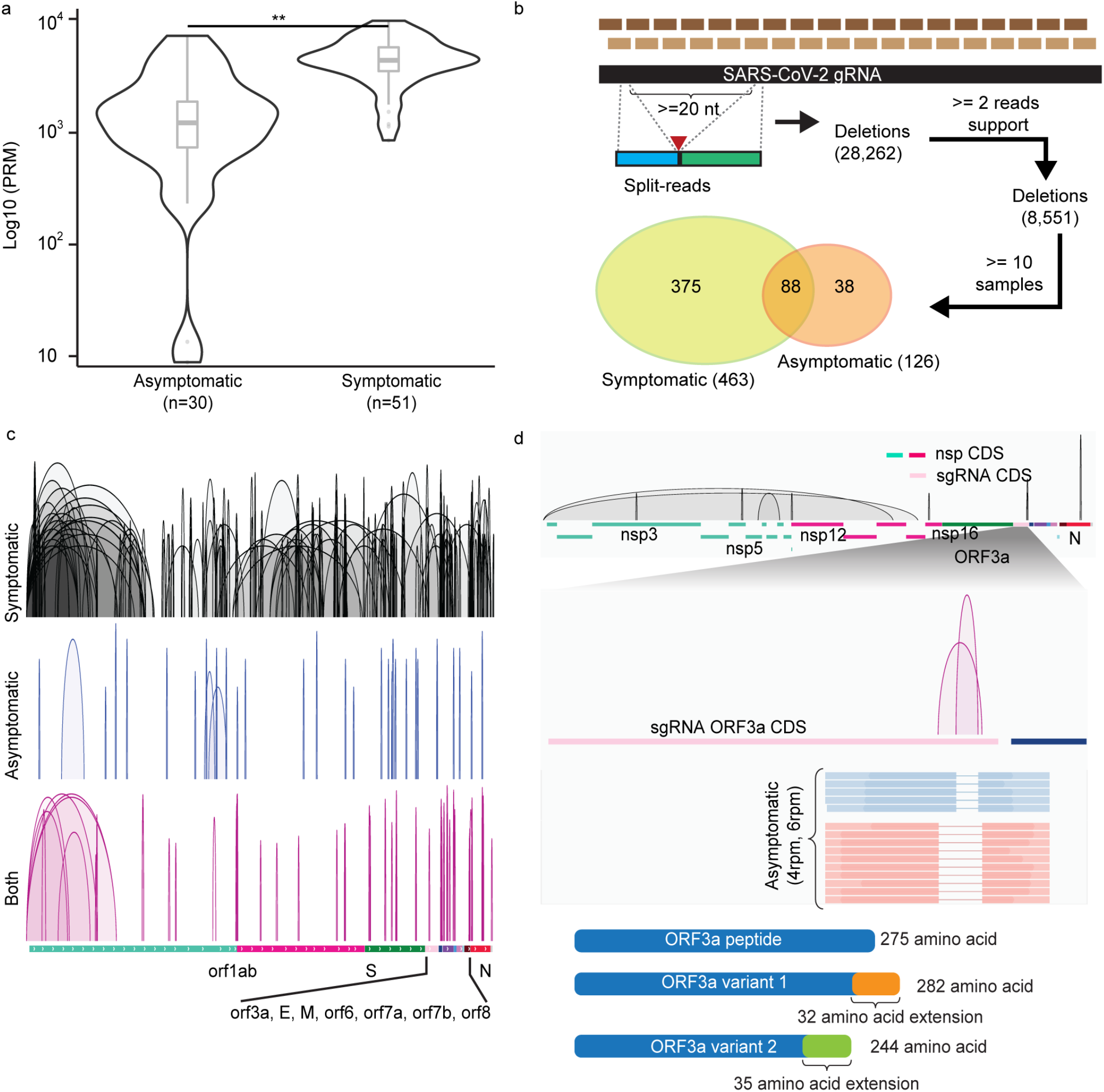
Deletions of SARS-CoV-2 RNAs in symptomatic and asymptomatic COVID-19 positive patients. **a.** Distributions of normalized split-aligned reads counts in asymptomatic and symptomatic patients. Two-sided Wilcoxon Rank-Sum Test, *p* = 2.3 × 10^−8^. Center line, median; boxes, first and third quartiles; whiskers, 1.5 × the interquartile range. **b.** Deletions inferred by amplicon-seq data from asymptomatic and symptomatic patients’ specimens. **c.** Visualization of the deletions detected in symptomatic (n=287), asymptomatic (n=34) and both (n=79) samples in IGV genome browser in reference annotated subgenomic RNA (sgRNA) transcribed regions. **d.** Top: Deletions (n=10) preferentially found in viral RNAs from the asymptomatic samples. Middle: zoom-in view in sgRNA_ORF3a coding sequence (CDS) region shows the two deletions uniquely found in asymptomatic cases, their normalized counts and representative read supports. Lower: their predicted translated peptide in reference to the wildtype ORF3a peptide.

These deletions were spread across the entire viral genome (Fig. 3c). To investigate the existence of distinct sets of deletions in viral RNAs selected in hosts with differences in disease severity, we examined their relative abundance (defined by normalized counts of read support) and frequencies (defined by the proportions of symptomatic vs asymptomatic samples found). We revealed 296 deletions significantly enriched in the symptomatic and 10 deletions in asymptomatic infections (*p*-value <0.05) (Supplementary Table 4). Among them, 263 and 9 deletions were exclusively found in symptomatic and asymptomatic specimens, respectively. We were particularly interested in the 10 deletions preferentially found in the asymptomatic hosts (Fig. 3d) and their impact on the integrity of viral sgRNAs and proteins. Notably, three of them located within the coding regions of sgRNAs and two of the three deletions (42 and 82 nucleotides, respectively) affected protein coding region of sgRNA_ORF3a. These deletions were predicted to yield ORF3a protein variants with C-terminal extension and truncation (Fig. 3d). ORF3a protein was shown to induce apoptosis in infected cells ^31^, an important host antiviral defense mechanism that controls the inflammatory response ^32^. The alteration in the ORF3a protein could weaken its pro-apoptotic activities which potentially reduce apoptosis-mediated immune responses and result in milder or even asymptomatic infection. Other asymptomatic-associated deletions were found within the coding regions of sgRNA_N, nsp5 and nsp16 which were predicted to yield truncations in encoded proteins for nucleocapsid, 3C-like proteinase and methyltransferase, respectively. Taken together, the existence of different types of deletions in viral RNAs exclusively observed in infected individuals exhibiting different host responses and their presence can be found in multiple independent infections strongly implicate the functional significance of structural variants in conferring features of SARS-CoV-2 virulence and pathogenicity.

### Full-length Iso-seq analysis revealed extensive structural variation in SARS-CoV-2 genomes

The widespread and abundant deletions arisen in the symptomatic infections drew our attention to investigate their diversity and impact on viral sgRNA transcription. The observed viral variants presumably resulted from deletions occurring either during viral replication or transcription (Fig. 4a). To distinguish their origins and characterize their impact on the viral translated protein products, we examined these deletions in the context of their associated sgRNA structures by full-length (FL) Iso-seq sequencing ^33^. From 10 samples with the highest ratio of SARS-CoV-2 content, we generated in total over two million of high-quality FL cDNA sequences (Supplementary Table 5). Of which, 632,207 (31%) of them were SAR-CoV-2 origins and were further clustered into 15,244 distinct transcript units (TUs) supported by ≥ 2 FL cDNA sequences (Fig. 4b). Based on their alignments across TRS-L and their respective canonical TRS-B junction sites, 1,114 FL TUs can be unambiguously assigned to sgRNA origins (Fig. 4c) while 4,591 FL TUs aligned uninterrupted across TRS-B site and were determined as products from viral gRNAs (Fig. 4b). When we examined the presence of deletions in these FL TUs, vast majority of the deletions were independently detected in both the sgRNA- and gRNA-derived FL TUs. Their validity was further supported by the breakpoints inferred from the split reads in the metatranscriptome RNA-seq data, suggesting that these were *bona fide* deletions occurred during viral gRNA replication as a result of low fidelity of RNA polymerases. These structural variants were subsequently propagated into protein-coding sgRNAs via transcription. Taking a TU of sgRNA_ORF3a as an example, this TU comprised 4 distinct deletions of 31, 34, 36 and 1,371 nucleotides, respectively which were independently uncovered by short-read RNA-seq data (Fig. 5a). The same deletions can be also found in in multiple TUs encoding distinct sgRNAs including sgRNA_E, _M and _ORF6 (Fig. 5b). Overall, from total of 15,244 FL TUs, 3,537 (23%) TUs harbored minimally one insertion or deletion over ≥20 bases, which raises the possibility that a substantial population of the SARS-CoV-2 virus carry structural variations during active infection. Therefore, structural variations of SARS-CoV-2 often lead to alternative sgRNA transcripts and significant alterations in their translation products. These variants potentially exist as quasispecies to facilitate evolutionary selection and host adaptation as observed in other RNA viral species ^34–37^.

**Figure 4:**
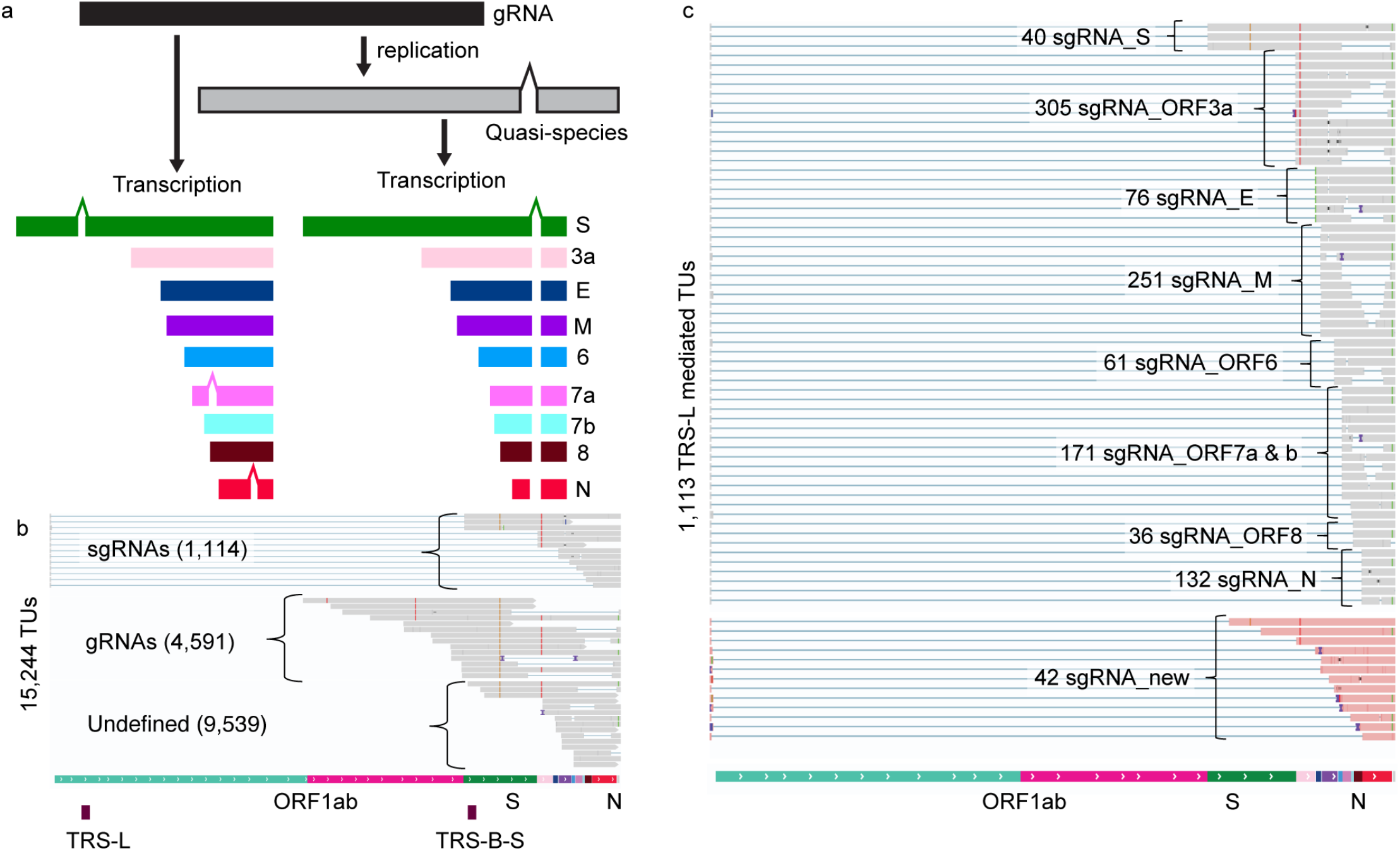
SARS-CoV-2 transcriptome diversity. **a.** Proposed models of the origins of SARS-CoV-2 genomic deletions resulted from the lack of accurate replications of viral gRNAs (Right) or transcription of viral sgRNA (Left). **b.** Assignment of **full-length (FL) transcript units (TUs)** revealed by long-read Iso-seq into viral sgRNAs (n=1,114), gRNAs (n=4,591) or undefined (n=9,539) based on their spans across Transcription regulatory sequence (TRS)-Leader/-Body (TRS-L/TRS-B) junctions. **c.** The distribution of FL TUs assigned to different sgRNA species based on their corresponding TRS-B sites.

**Figure 5.**
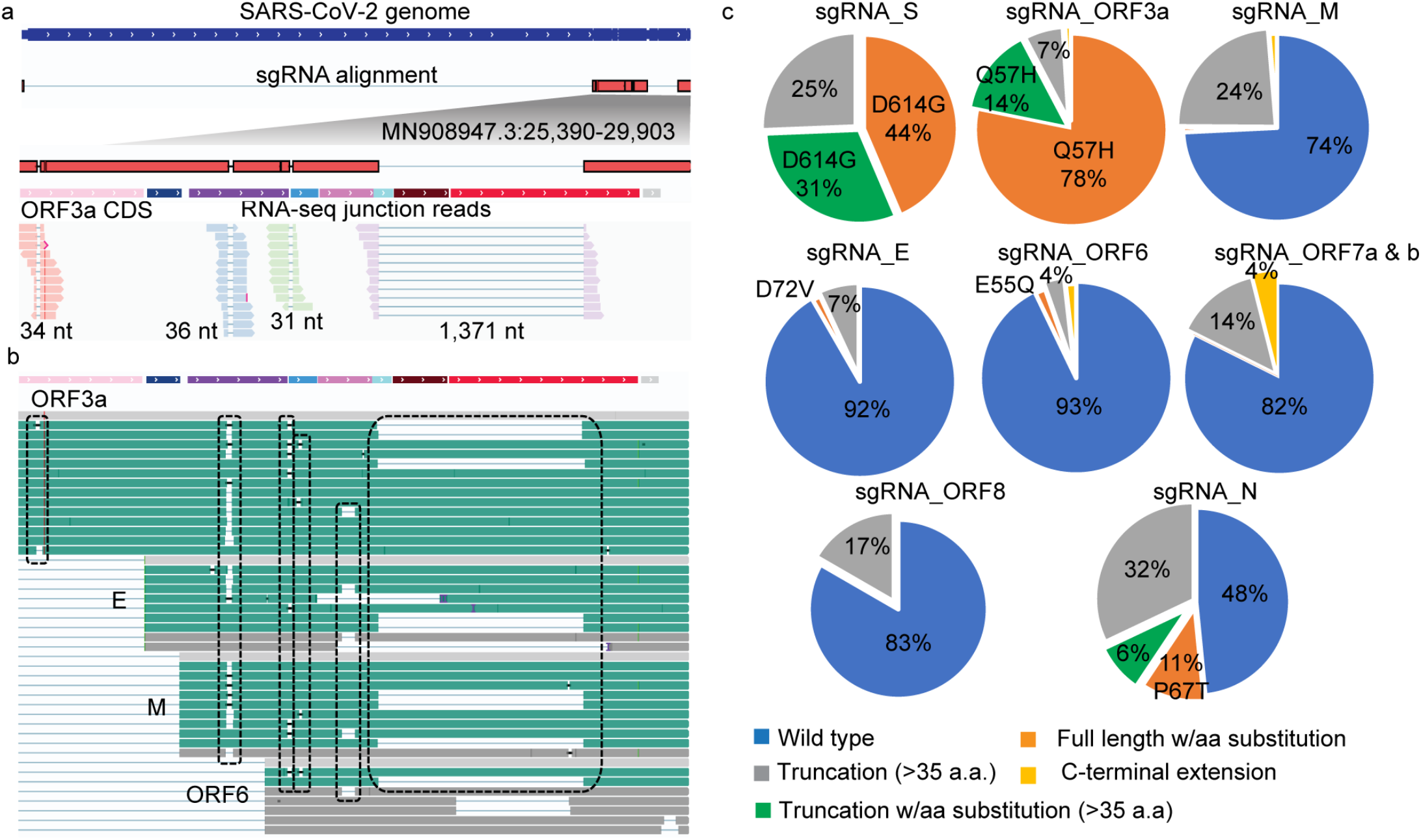
Phasing structural variants on subgenomic RNA (sgRNA) derived full-length (FL) transcript units (TUs) reveals SARS-CoV-2 proteome complexity. **a.** An example of 4 deletions co-occurred on a sgRNA_ORF3a molecule uncovered by FL iso-seq analysis. **b.** Examples show identical deletions (highlighted in boxes) detected in multiple FL cDNAs encoding different sgRNA species. **c.** Distribution of predicted wild-type and mutant proteins encoded from the sgRNA derived FL TUs.

### Structural variants in viral genomes further expand viral proteome complexity

Through placing the co-occurred insertions and deletions onto the individual FL transcripts, we can investigate the precise impacts of these variants on the viral protein translation. From the collection of the 1,114 sgRNA-derived FL cDNA sequences, 23% of these transcripts carrying frameshifts with >35aa predicted translated protein products of truncations (20.1%), extension (1.2%) and new peptides of no known functional annotation (1.3%). Intriguingly, we also observed low frequency of FL cDNAs producing potential fusion proteins. For example, a 257 amino-acid Membrane and ORF6 fusion peptide resulted from a 31-bases deletion. From the combinatorial effects of the non-synonymous SNVs and detected indels, the diversity of the SARS-CoV-2 encoded proteome were derived for each of the sgRNA species. We found the translated proteins as the following five groups: 1) Wild type proteins of known annotation. 2) Proteins of known annotation with amino acid substitutions. 3) Truncated proteins of known annotation with or without amino acid substitutions. 4) Proteins of known annotation with C-terminal extension, and 5) New peptides. The proportions of the wild-type proteins and their corresponding variant types for each of the eight sgRNA-encoded proteins were shown in Fig. 5c. As expected, vast majority of the predicted structural and accessory proteins translated from sgRNAs detected in these clinical samples were full-length forms. Among the eight sgRNA-encoded proteins, ORF6 and Envelop are the most stable with 93% and 92% predicted FL wild-type proteins.

The predominant forms of S and ORF3a carry amino acid substitutions D614G and Q57H resulted from the non-synonymous SNVs in MN908947.3:25563 (G>U) and MN908947.3:23403 (A>G), respectively. SARS-CoV-2 D614G variant, emerging early during the pandemic, was suggested to possess higher infectivity ^38^ while the effect of Q57H variant on viral pathophysiology is currently less clear. Similar to D614G, Q57H variant could be subjected to natural selection because it was only reported at < 6% in Feb 2020 ^39^. 56% of Spike and 41% of Nucleocapsid were predicted to be truncated. We further annotated the deleted regions for function domains using the NCBI conserved domain database (CDD) ^40^ and, to our surprise, we found that 41% and 42% of the predicted truncated Spike and Nucleocapsid proteins were lacking the receptor-binding domain (RBD) and RNA-binding domain (PSSM-ID 394862), respectively. S protein functions to mediate host cell entry through angiotensin-converting enzyme 2 (ACE2) receptor binding ^41^ and RNA-binding domain in N protein plays an important role in virus transcription and assembly ^42^. These proteins are widely used as targets for vaccine and drug development ^43^, with some exclusively targeting the RBD for treatment with neutralizing antibodies ^44^. While the high frequencies of structural deletion in these proteins were only observed in selected samples with high viral content, if verified in a larger population of the infected human cells, they could have significant ramifications on the efficacy of antibody-induced immunity and devising treatment strategies.

## Discussion

In this study, we attempted to study the activity of SARS-CoV-2 transcription and the complexity of viral genome structural variation in infected human hosts with distinct disease severity. Through a combination of multi-scale genomic analyses, we quantitatively evaluated the expression of sgRNA species in a broad range of swabs collected for routine PCR-based diagnostics and revealed that the relative abundance of sgRNAs were significantly lower in the infected individuals without COVID-19 associated symptoms, indicating repressed viral transcription. The lower levels of sgRNAs detected in the asymptomatic infection was unlikely due to the timing of the sample collections, i.e. pre-symptomatic because sgRNAs are thought to be abundant in early infection ^17^. Moreover, the repression of sgRNA was not attributed to the differences in viral load and the sgRNAs quantities were normalized with the levels of gRNAs in each sample.

Different from diagnostic RT-qPCR assays which mainly measure the viral genomic RNA shedding, characterizing viral sgRNAs in the COVID-19 positive samples could be informative to understand the virus’ replicative activity in the host cells. Previous studies showed an increase of viral load is indicative of an aggravation of symptoms ^17^ and the detection of sgRNAs also positively correlated with the isolation of infectious virus in tissue cultures ^45^. Building from these observations, our results further show that sgRNA levels as assessed by the sgRNA/gRNA ratio are highly correlated with one measure of clinical severity, the presence of symptoms. The more rapid viral clearance seen in asymptomatic patients may result from successful host immune responses. Our sgRNA findings suggest that RT-qPCR based assays to quantitatively evaluate the relative abundance of sgRNAs may be a predictive measure of the clinical severity of COVID-19 symptoms. These results could have significant impact on conservation of medical resources during the rapid community spreading, much like what we are experiencing globally in recent weeks.

Our work also showed distinct and recurring sets of viral RNA deletions in both symptomatic and asymptomatic infections. Their consistent and preferential detection in multiple COVID-19 positive cases point to the genome instability as a source of viral proteome complexity and potential evolutionary selection for host adaptation. Taken together, when associated together with the host genetics and immune response, the sgRNA expression and structural diversity could provide insight in understanding host-viral interactions, evolution and transmission. This, in turn, will guide risk mitigation, testing strategies, and inform future vaccine development.

## Methods

### Sample Collection

Samples for the clinical diagnosis purpose were collected by a combination of nasal, oral, nasopharyngeal and oropharyngeal swabs between April to September 2020. Patient age ranged from 18 to 97 years (median 67 years); 35 were male and 45 female (ST. 1). Specimens collected were swabs of nasopharyngeal (n=42), anterior nasal (n=35), and oropharyngeal (n=5). The swabs preserved in viral transport media that were kept at 4 - 8 °C for less than 72 hours between collection and testing. Among them, 51 of the samples were collected from patients presented at the hospital with symptoms consistent with COVID-19 and 30 of the samples were collected from the screening programs. All 81 underwent testing through RT-PCR by TaqPath™ COVID-19 Combo Kit (ThermoFisher) under the FDA Emergency Use Authorization (EUA) with confirmed positive diagnosis.

### Amplicon sequencing and data processing

Total RNA was extracted from 81 clinical COVID-19 confirmed positive samples using the MagMAX Viral/Pathogen Nucleic Acid Isolation Kit on the KingFisher Flex. The extracted RNAs were used for first strand cDNA synthesis priming with random hexamer using SuperScript IV as per manufacturers’ instructions. The cDNAs were amplified in two multiplex PCR reactions using the multiplex PCR primers (V3) tiled across the viral genome developed by the ARCTIC Network ^46^ to PCR-amplify the viral genome with primers. The amplicons were pooled and ligated with Illumina UDI adaptor (Illumina). Product were PCR amplified by 5 cycles and cleaned up using SPRI beads (Beckman Coulter) and subjected to paired-end 300bp sequencing on Illumina Miseq. Raw paired-end reads were trimmed with trim_galore [https://github.com/FelixKrueger/TrimGalore] (v0.4.3) via cutadapt ^47^ (v1.2.1) with the parameters “--stringency 3 −q 30 −e ․10 --length 15 --paired”. The trimmed reads were classified with centrifuge-1.0.3-beta ^28^ for their potential source. They were aligned to the SAR-Cov2 reference (MN908947.3) with STAR ^48^ (v2.7.3a) with many switches to completely turn off the penalties of non-canonical eukaryotic splicing as documented ^21^: “--outFilterType BySJout --outFilterMultimapNmax 20 --alignSJoverhangMin 8 --alignSJDBoverhangMin 1 --outSJfilterOverhangMin 12 12 12 12 --outSJfilterCountUniqueMin 1 1 1 1 -- outSJfilterCountTotalMin 1 1 1 1 --outSJfilterDistToOtherSJmin 0 0 0 0 -- outFilterMismatchNmax 999 --outFilterMismatchNoverReadLmax 0.04 --scoreGapNoncan −4 --scoreGapATAC −4 --chimOutType Junctions WithinBAM HardClip --chimScoreJunctionNonGTAG 0 --alignSJstitchMismatchNmax −1 −1 −1 −1 --alignIntronMin 20 --alignIntronMax 1000000 --alignMatesGapMax 1000000”. We retained aligned paired-end reads which start with the primer-binding site mutually exclusive from the primers Pool 1 or Pool 2 at the 5’ end of both R/1 and R/2. These retained paired end reads CIGAR was parsed for jumps and deletions (represented by CIGAR operations N or D of size ≥20 bases).

### SARS-CoV-2 sgRNAs and gRNA expression in the Amplicon-seq data

The TRS-L site is located in amplicon 1 of primers Pool 1. Thus, only sgRNAs with TRS-B sites present in the amplicons from primers Pool 1 can be detected. The six detectable sgRNAs are sgRNA_S (Primers 1-and-71), sgRNA_E (Primers 1-and-87), sgRNA_M (Primers 1-and-87), sgRNA_6 (Primers 1-and-89_alt2), sgRNA_7b (Primers 1-and-91), and sgRNA_N (Primers 1- and-93). To classify an aligned paired-end read as originated from sgRNA, it must contain the mentioned primers binding sites from one of the six detectable sgRNAs. Additionally, it must contain at least one split-aligned read thats split read junction marks the leader-to-body junction and that the translated protein product from the concatenated sequence produces the canonical sgRNA. The rest of the amplicon 1 aligned pair-end reads are classified as originated from gRNA.

All sgRNA expression is inter-sample normalized by a scale factor of 1,000,000/ total number of mapped read-pairs, giving a comparable measure unit read-pair per million (RPM). The ratio of sgRNA/gRNA is simply computed as the ratio of aligned read-pairs in amplicon 1 as follow: the number of split-aligned read-pairs covering the genomic position 31-75 to the number of read-pairs covering the genomic position 31-410 without split-alignment.

### Short-read RNA sequencing and data processing

RNA-seq libraries were prepared with KAPA mRNA HyperPrep Kit (Roche) according to manufacturer’s instruction. First, poly-A^+^ RNA was isolated from 1ul of total RNA extracted from clinical samples using oligo-dT magnetic beads. Purified RNA was then fragmented at 85°C for 6 mins, targeting fragments range 250-300bp. Fragmented RNA is reverse transcribed with an incubation of 25°C for 10mins, 42°C for 15mins and an inactivation step at 70C for 15mins. This was followed by second strand synthesis and A-tailing at 16°C for 30mins and 62°C for 10min. A-tailed, double stranded cDNA fragments were ligated with Illumina-compatible adaptors with Unique Molecular Identifier (UMI) (IDT). Adaptor-ligated DNA was purified using Ampure XP beads (Beckman Coultier). This is followed by 17 cycles of PCR amplification. The final library was cleaned up using AMpure XP beads. Quantification of libraries were performed using real-time qPCR (Thermo Fisher). Sequencing was performed on Illumina Novaseq paired-end 149 bases with indexes and 9 bases of UMI. Raw paired-end reads were trimmed, potential source classified, and mapped per documented above (Amplicon data processing). Reads deduplication were performed with UMI-tools (v1.0.1) ^49^. The aligned paired end reads CIGAR was parsed for jumps and deletions (represented by CIGAR operations N or D of size ≥20 bases).

### Viral load vs sgRNA abundance

Samples with ≥100 UMI-deduplicated split-aligned read-pairs are considered (n=45). The sgRNA abundance inter-sample normalized by a scale factor of 1,000,000/total number of UMI-deduplicated mapped read-pairs, giving a comparable measure unit (junction-)read-pair per million (RPM) The sample viral load is calculated by transforming the *Ct* value with 2 to the power of (27-*Ct*). The value 27 is chosen to allow calculated values to be comparable to the numbers of junction-read per million reads.

### Define genomic RNA and canonical sgRNA reads from Illumina RNA-seq data

We followed ^21^ definition of read classification for sgRNA with a modification. We still required that the split read junction to mark the leader-to-body junction and that the translated protein product from the concatenated sequence produces the canonical sgRNA. However, we require that split read 5’ site of deletion is mapped to a genomic position between 59 and 79 (TRS-L: 70-75 nt), instead of 55 and 85 ^21^. This is established based on the sequence identity between the leader and body regions. For comparable gRNA read count (with respect to sgRNAs read counts), we require that the read must harbor no junction, must overlap the genomic position 1 to 85, and its mate read must mapped within the first 1000 base of the genome.

The relative abundance of a sample’s sgRNA is, thus, the sgRNA read counts over the sum of the sample’s gRNA and all sgRNAs read count.

### Genomic RNA and canonical sgRNA abundance in Vero cell

DNBseq RNA sequencing data of SARS-CoV-2-infected Vero cell ^21^ was downloaded. The data was processed, and expression computed exactly per our short-read RNA sequencing data.

### Long-read Iso-seq and data processing

Total RNA extracted from nasopharyngeal swabs were prepared according to Iso-seq Express Template Preparation (Pacbio). Full length cDNA is generated using NEBNext Single Cell/ Low Input cDNA synthesis and Amplification Module in combination with Iso-seq Express Oligo Kit. Amplified cDNA is purified using ProNex beads. For samples with lower than 160ng in yield, additional PCR cycles is added. cDNA yield of 160ng-500ng were then underwent SMRTbell library preparation including a DNA damage repair, end repair and A-tailing and finally ligated with Overhang Barcoded Adaptors. Libraries were then pooled and sequenced on Pacbio Sequel II. The raw sequencing data generated were processed with the SMRT Link (v 8.0.0.80529) Iso-Seq analysis pipeline with the default parameters. Firstly, circular consensus sequences (CCSs) were generated from the raw sequencing reads. Demultiplexed CCSs based on sample barcodes in the adaptors, were further classified into full length, non-chimeric (FLNC) CCSs and non-full length, non-chimeric CCSs based on the presence of chimera sequence, sequencing primer and 3’ terminal poly-A sequence. FLNC CCSs (which contains both the 5’-and-3’-adaptor sequence along with the poly-A tail) were clustered to generate isoforms. Only the high-quality (accuracy≥0.99) transcript isoforms (referred here as TUs) were aligned to the SARS-CoV-2 genome reference (MN908947.3) with pbmm2 (v1.1.0). The aligned TU’s CIGAR was parsed for gaps (represented by CIGAR operations N or D of size ≥20bases). The identified gaps were clustered based on their aligned genomic coordinates. The maximum difference amongst the cluster members’ gap start (and end) coordinates is 10 bases. For TU with multiple transcribed segments, and its first segment 3’ site mapped to the genomic position 59-79, the TU is considered TRS-L mediated. The translation products of the TUs were predicted by translating the sequence with standard genetic code upon the first AUG (Methionine) encountered. The translation product is annotated against Conserved Domain Database (CDD) including 55,570 position-specific score matrices (PSSMs) ^40^.

## Supporting information

Supplementary Table 1. Sample summary

Supplementary Table 2. Amplicon-Seq data summary

Supplementary Table 3. Poly-A+ RNA-Seq data summary

Supplementary Table 4A. Enriched mutations

Supplementary Table 5. Iso-Seq data summary

## Data availability statement

All data described in this study has been deposited in NCBI’s Sequence Read Archive PRJNA690577. (https://www.ncbi.nlm.nih.gov/bioproject/690577)

## Acknowledgements

The authors thank all Jackson Laboratory Clinical Laboratory team members for their effort in samples collection and processing of covid-19 samples; and the Jackson Laboratory Genome Technologies team members for their sequencing effort. We also thank Linda Choquette and her Clinical & Translational Research Support team on coordination effort in patient clinical information collection. Research reported in this publication was supported by The Jackson Laboratory Scientific Service Innovation Fund (JAX-SSIF-FY20-CLW-SARS-CoV-2) awarded to C.-L.W. C.-L.W. and C.Y.N. are supported by NCI under Award Number P30CA034196.

## Reference

1. Buitrago-Garcia, D. et al. Occurrence and transmission potential of asymptomatic and presymptomatic SARS-CoV-2 infections: A living systematic review and meta-analysis. PLoS Med 17, e1003346 (2020).

2. Byambasuren, O. et al. Estimating the extent of asymptomatic COVID-19 and its potential for community transmission: Systematic review and meta-analysis. Official Journal of the Association of Medical Microbiology and Infectious Disease Canada, e20200030 (2020).

3. Ing, A.J., Cocks, C. & Green, J.P. COVID-19: in the footsteps of Ernest Shackleton. Thorax 75, 693–694 (2020).

4. Xiao, T. et al. Early viral clearance and antibody kinetics of COVID-19 among asymptomatic carriers. medRxiv, 2020.04.28.20083139 (2020).

5. Chau, N.V.V. et al. The natural history and transmission potential of asymptomatic SARS-CoV-2 infection. Clin Infect Dis (2020).

6. Hu, Z. et al. Clinical characteristics of 24 asymptomatic infections with COVID-19 screened among close contacts in Nanjing, China. Sci China Life Sci 63, 706–711 (2020).

7. Yang, R., Gui, X. & Xiong, Y. Comparison of Clinical Characteristics of Patients with Asymptomatic vs Symptomatic Coronavirus Disease 2019 in Wuhan, China. JAMA Netw Open 3, e2010182 (2020).

8. Hurst, J.H. et al. SARS-CoV-2 Infections Among Children in the Biospecimens from Respiratory Virus-Exposed Kids (BRAVE Kids) Study. medRxiv (2020).

9. Nogrady, B. What the data say about asymptomatic COVID infections. Nature 587, 534–535 (2020).

10. Lavezzo, E. et al. Suppression of a SARS-CoV-2 outbreak in the Italian municipality of Vo’. Nature 584, 425–429 (2020).

11. Arons, M.M. et al. Presymptomatic SARS-CoV-2 Infections and Transmission in a Skilled Nursing Facility. N Engl J Med 382, 2081–2090 (2020).

12. Alm, E. et al. Geographical and temporal distribution of SARS-CoV-2 clades in the WHO European Region, January to June 2020. Euro Surveill 25(2020).

13. Hadfield, J. et al. Nextstrain: real-time tracking of pathogen evolution. Bioinformatics 34, 4121–4123 (2018).

14. https://nextstrain.org/sars-cov-2/

15. Sola, I., Almazan, F., Zuniga, S. & Enjuanes, L. Continuous and Discontinuous RNA Synthesis in Coronaviruses. Annu Rev Virol 2, 265–88 (2015).

16. Cui, J., Li, F. & Shi, Z.L. Origin and evolution of pathogenic coronaviruses. Nat Rev Microbiol 17, 181–192 (2019).

17. Wolfel, R. et al. Virological assessment of hospitalized patients with COVID-2019. Nature 581, 465–469 (2020).

18. de Haan, C.A., Masters, P.S., Shen, X., Weiss, S. & Rottier, P.J. The group-specific murine coronavirus genes are not essential, but their deletion, by reverse genetics, is attenuating in the natural host. Virology 296, 177–89 (2002).

19. Yount, B. et al. Severe acute respiratory syndrome coronavirus group-specific open reading frames encode nonessential functions for replication in cell cultures and mice. J Virol 79, 14909–22 (2005).

20. Davidson, A.D. et al. Characterisation of the transcriptome and proteome of SARS-CoV-2 reveals a cell passage induced in-frame deletion of the furin-like cleavage site from the spike glycoprotein. Genome Med 12, 68 (2020).

21. Kim, D. et al. The Architecture of SARS-CoV-2 Transcriptome. Cell 181, 914–921 e10 (2020).

22. Nomburg, J., Meyerson, M. & DeCaprio, J.A. Pervasive generation of non-canonical subgenomic RNAs by SARS-CoV-2. Genome Med 12, 108 (2020).

23. Young, B.E. et al. Effects of a major deletion in the SARS-CoV-2 genome on the severity of infection and the inflammatory response: an observational cohort study. Lancet 396, 603–611 (2020).

24. https://www.gov.uk/government/publications/investigation-of-novel-sars-cov-2-variant-variant-of-concern-20201201

25. Tyson, J.R. et al. Improvements to the ARTIC multiplex PCR method for SARS-CoV-2 genome sequencing using nanopore. bioRxiv (2020).

26. Gohl, D.M. et al. A rapid, cost-effective tailed amplicon method for sequencing SARS-CoV-2. BMC Genomics 21, 863 (2020).

27. MacKay, M.J. et al. The COVID-19 XPRIZE and the need for scalable, fast, and widespread testing. Nat Biotechnol 38, 1021–1024 (2020).

28. Kim, D., Song, L., Breitwieser, F.P. & Salzberg, S.L. Centrifuge: rapid and sensitive classification of metagenomic sequences. Genome Res 26, 1721–1729 (2016).

29. https://www.cdc.gov/coronavirus/2019-ncov/more/scientific-brief-emerging-variant.html

30. https://virological.org/t/preliminary-genomic-characterisation-of-an-emergent-sars-cov-2-lineage-in-the-uk-defined-by-a-novel-set-of-spike-mutations/563

31. Ren, Y. et al. The ORF3a protein of SARS-CoV-2 induces apoptosis in cells. Cell Mol Immunol 17, 881–883 (2020).

32. Roulston, A., Marcellus, R.C. & Branton, P.E. Viruses and apoptosis. Annu Rev Microbiol 53, 577–628 (1999).

33. Wang, B. et al. Unveiling the complexity of the maize transcriptome by single-molecule long-read sequencing. Nat Commun 7, 11708 (2016).

34. Xue, K.S., Hooper, K.A., Ollodart, A.R., Dingens, A.S. & Bloom, J.D. Cooperation between distinct viral variants promotes growth of H3N2 influenza in cell culture. Elife 5, e13974 (2016).

35. Domingo, E., Sheldon, J. & Perales, C. Viral quasispecies evolution. Microbiol Mol Biol Rev 76, 159–216 (2012).

36. Chaudhry, M.Z. et al. SARS-CoV-2 Quasispecies Mediate Rapid Virus Evolution and Adaptation. bioRxiv, 2020.08.10.241414 (2020).

37. Jary, A. et al. Evolution of viral quasispecies during SARS-CoV-2 infection. Clin Microbiol Infect 26, 1560 e1–1560 e4 (2020).

38. Korber, B. et al. Tracking Changes in SARS-CoV-2 Spike: Evidence that D614G Increases Infectivity of the COVID-19 Virus. Cell 182, 812–827 e19 (2020).

39. Koyama, T., Platt, D. & Parida, L. Variant analysis of SARS-CoV-2 genomes. Bull World Health Organ 98, 495–504 (2020).

40. Lu, S. et al. CDD/SPARCLE: the conserved domain database in 2020. Nucleic Acids Res 48, D265–D268 (2020).

41. Letko, M., Marzi, A. & Munster, V. Functional assessment of cell entry and receptor usage for SARS-CoV-2 and other lineage B betacoronaviruses. Nat Microbiol 5, 562–569 (2020).

42. McBride, R., van Zyl, M. & Fielding, B.C. The coronavirus nucleocapsid is a multifunctional protein. Viruses 6, 2991–3018 (2014).

43. Ahmed, S.F., Quadeer, A.A. & McKay, M.R. Preliminary Identification of Potential Vaccine Targets for the COVID-19 Coronavirus (SARS-CoV-2) Based on SARS-CoV Immunological Studies. Viruses 12(2020).

44. Salvatori, G. et al. SARS-CoV-2 SPIKE PROTEIN: an optimal immunological target for vaccines. J Transl Med 18, 222 (2020).

45. Perera, R. et al. SARS-CoV-2 Virus Culture and Subgenomic RNA for Respiratory Specimens from Patients with Mild Coronavirus Disease. Emerg Infect Dis 26, 2701–2704 (2020).

46. https://www.protocols.io/view/ncov-2019-sequencing-protocol-v3-locost-bh42j8ye

47. Martin, M. Cutadapt removes adapter sequences from high-throughput sequencing reads. EMBnet.journal; Vol 17, No 1: Next Generation Sequencing Data Analysis (2011).

48. Dobin, A. et al. STAR: ultrafast universal RNA-seq aligner. Bioinformatics 29, 15–21 (2013).

49. Smith, T., Heger, A. & Sudbery, I. UMI-tools: modeling sequencing errors in Unique Molecular Identifiers to improve quantification accuracy. Genome Res 27, 491–499 (2017).

